# Bayesian Inference of Two-Dimensional Contrast Sensitivity Functions

**DOI:** 10.1101/067116

**Authors:** Xiaoxiao Wang, Huan Wang, Jinfeng Huang, Tzvetomir Tzvetanov, Yifeng Zhou

## Abstract

The contrast sensitivity function (CSF) is crucial in predicting functional vision both in research and clinical areas. Recently, a group of novel strategies, multi-dimensional adaptive methods, were proposed and allowed more rapid measurements when compared to usual methods such as Ψ or staircase. Our study further presents a multi-dimensional Bayesian framework to estimate parameters of the CSF from experimental data obtained by classical sampling. We extensively simulated the framework’s performance as well as validated the results in a psychophysical experiment. The results showed that the Bayesian framework significantly improves the accuracy and precision of parameter estimates from usual strategies, and requires about the same number of observations as the novel methods to obtain reliable inferences. Additionally, the improvement with the Bayesian framework was maintained when the prior poorly matched the observer’s CSFs. The results indicated that the Bayesian framework is flexible and sufficiently precise for estimating CSFs.

## Introduction

Vision science, whose aim is to provide a mechanistic explanation of human vision, has placed a great importance in measuring and explaining contrast sensitivity over a wide range of spatial frequencies (Pelli & Bex, 2013). The spatial contrast sensitivity function (CSF), which relates subject’s response to stimulus spatial frequency (SF) and contrast, is accepted as a basic comprehensive measure of the visual system in both normal and abnormal vision (Ginsburg, 2003; Hess, France, & Tulunay-Keesey, 1981; Jindra & Zemon, 1989; Regan, Raymond, Ginsburg, & Murray, 1981), and is one of the most important metrics in investigating functional deficits in visual disorders (Hess & Howell, 1977; Hot, Dul, & Swanson, 2008; Hou et al., 2010; C.-B. Huang, Zhou, & Lu, 2008; C. Huang, Tao, Zhou, & Lu, 2007; Zhou et al., 2006). However, the CSF, that spans two feature dimensions (contrast×SF), has the disadvantage of taking a long time to sample.

Recently, it has been demonstrated that the CSF could be treated as a two-dimensional (2-D) psychometric function (Figure 1) and can be efficiently sampled by 2-D adaptive strategies. Based on a parametric model, these fancy methods search the 2-D stimulus space for the next most informative stimulus, allowing a more efficient estimation of the threshold contour than usual procedures (Hou et al., 2010; Kujala & Lukka, 2006; L. A. Lesmes, Lu, Baek, & Albright, 2010). These usual procedures, such as QUEST, Ψ, and staircases, maps responses to only one dimension (the contrast) of stimulus and need repetitive measurements along the other feature (the SF), which seems inefficient. At the same time, they have advantages of fairly simplicity in algorithm and fewer assumptions of function shape, and are widely applied by researchers in recent years (Bonneh, Adini, & Polat, 2016; Chung & Legge, 2016; Klein, 2001; Richard, Johnson, Thompson, & Hansen, 2015; Vedamurthy, Nahum, Bavelier, & Levi, 2015). An intriguing question is: could these simple usual methods be efficient, at least comparable with the fancy 2-D parametric adaptive methods, if Bayesian inference is also applied to them? The answer to this question could be extremely important for clinical application and may have important implication for the development of quicker inference protocols for CSFs.

**Figure 1.**
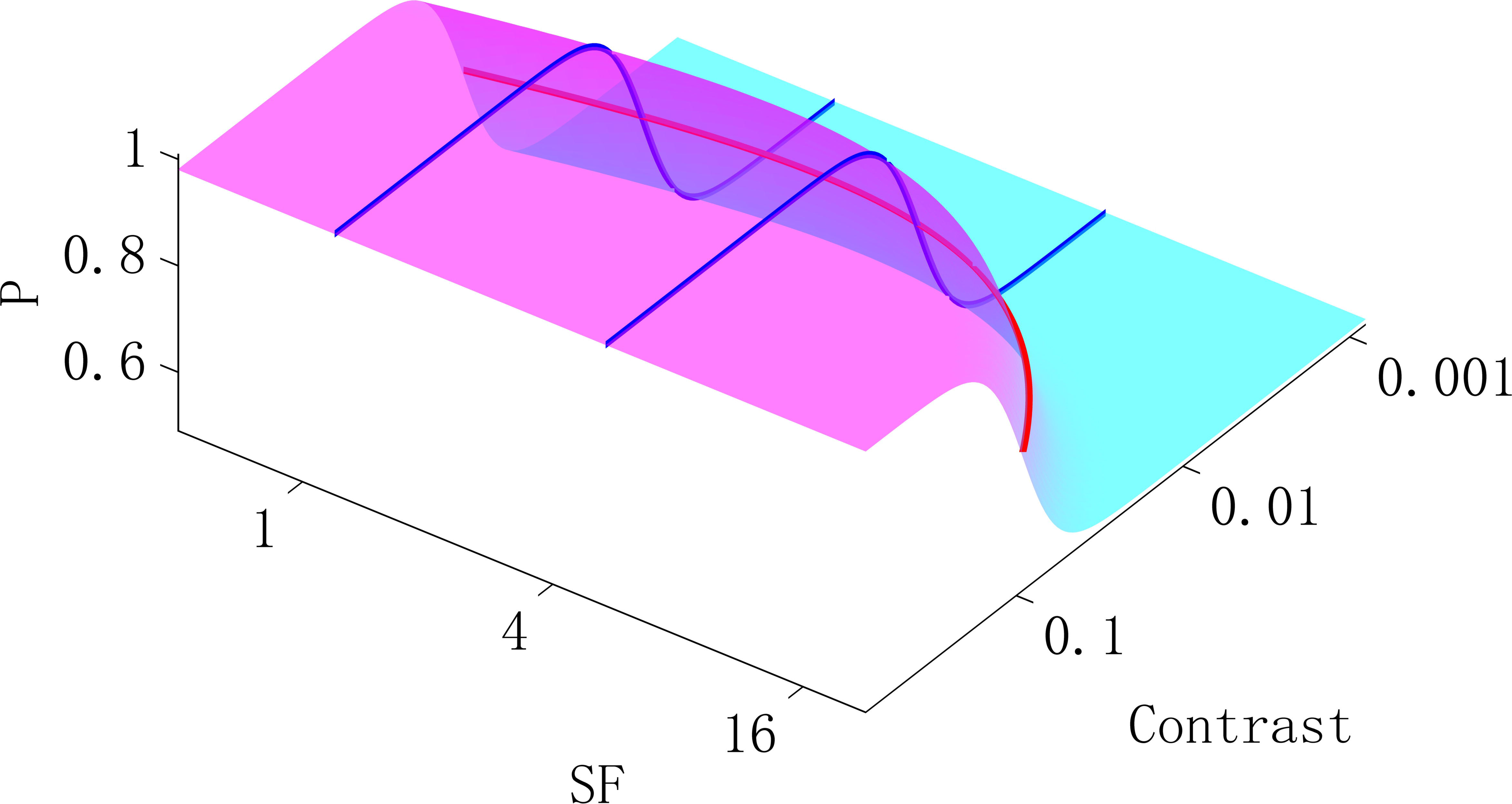
Illustration of 2-D spatial contrast sensitivity psychometric function. We use the reciprocal of the standard 1D CSF function across spatial frequencies (red curve) as midpoint in a logistic psychometric function along the contrast dimension (blue curve) and thus define the 2-D psychometric function.

In this study, we introduced a Bayesian framework for estimating CSF from experiments sampled by usual psychometric methods. We parameterized the 2D CSF with a logistic psychometric function along the log-contrast dimension together with a double-exponential for the SF dimension (Movshon & Kiorpes, 1988) (Figure 1). Given a set of observed data, the framework kept updating on a trial basis the posterior probability distribution of the parameters following Bayes’ theorem (Watson & Pelli, 1983). The final estimate of parameters is given by the mean of this posterior distribution (Kontsevich & Tyler, 1999).

We also assessed the Bayesian framework’s reliability and efficiency. Four adaptive procedures for next stimulus selection were evaluated: (1) two simple adaptive strategies – the simple staircase method (Kaernbach, 1991) which steps ‘up’ or ‘down’ the stimulus intensity after every ‘negative’ or ‘positive’ response respectively, and the Ψ method (Kontsevich & Tyler, 1999) which uses parametric adaptive techniques to place the next stimulus to minimize the expected entropy of the threshold and slope along the contrast dimension; (2) two novel 2-D adaptive methods: the qCSF (L. A. Lesmes et al., 2010) that optimized the sampling along the entire CSF curve and searches for the stimulus that minimized the expected entropy in both contrast and SF spaces, and the FIG (Fisher information gain) method (Remus & Collins, 2007), here adapted to 2D, that selects the next 2-D stimulus which maximizes the Fisher information gain of the function parameters. While assessing, the model of CSF, the levels of spatial frequency measured, the levels of contrast, and the amount of sampling trials were kept identical across these methods. The results allowed us to reveal, by the 2-D Bayesian framework, how much improvement of efficiency is available to the usual strategies.

## 1. Material and Methods

### 1.1. 2-D Psychometric Function

The usual psychometric function along contrast dimension is a one-dimensional (1-D) function Ψ_¸_(x), that represents the probability of the subject to detect a stimulus intensity x. This function is commonly chosen to have a sigmoid form and here we choose the Logistic function. The parameters are (α, β, γ, σ), where α denotes threshold, β is the slope, and the asymptotes γ and δ specify the guessing and lapsing rate. For an n-alternative forced choice (nAFC) task, where the subject is asked to choose between n possibilities, the value of γ should be equal to 1/n. Here, we restrict ourselves to the most common one, the standard 2AFC paradigm (γ =1/2).

The contrast sensitivity function (CSF), S(f), in its basic 1D representation describes the sensitivity (1/threshold) as a function of grating frequency (Wilson & Wilkinson, 2003). Here we use the double-exponential form (Movshon & Kiorpes, 1988; Figure 1, red curve), to describe the CSF:

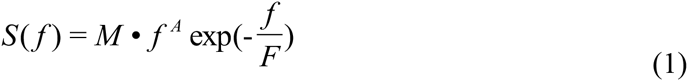

The parameters A and F relate to the steepness of the low- and high-frequency portions of the curve, F·A defines the peak spatial frequency and M(FA)^A^ exp(-A) its amplitude. We use it as midpoint in a logistic psychometric function and assume that the slope parameter does not vary with spatial frequency (Mayer & Tyler, 1986). Thus we define the 2-D log-log contrast-SF psychometric function as:

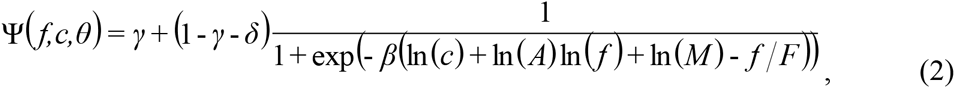

Where Ψ(f,c,θ) is the probability of ‘detect’ response at intensity x = (f, c) for the given parameters θ=(M, A, F, β, γ, δ).

### 1.2. Bayesian Inference

We used the Bayesian rule described in details by Kuss, Jäkel, and Wichmann (2005) and Kontsevich and Tyler, (1999). In a Bayesian psychometric function inference, a prior distribution is first assumed, representing beliefs about the value of the true parameters before the inference. Given the observed data, Bayes’ rule yields the posterior distribution:

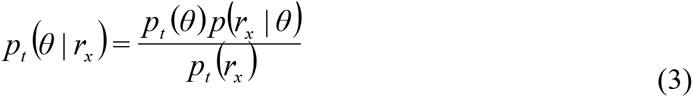

where the normalizing constant is given by: 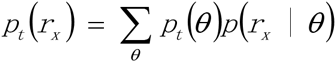, r_x_ denoting the response to the trial whose stimulus intensity is x. After each trial, the posterior distribution becomes the prior distribution of the next trial. In the experimental studies, the data set collected D = {(x,r_x_)_t_| t = 1, …, k} allows to sequentially update the posterior distribution on a trial basis, and thus to learn or improve the estimates of parameter p(θ | r_x_). The posterior distribution after all trials (Figure 6, magenta curve) is used to have a final estimate of the parameters θ. For the 2-D CSF psychometric function (Equation 2), γ is fixed at 0.5 according to the 2AFC design, and for any n-alternative forced choice (nAFC) task, γ should be fixed at 1/n.

Ideally, a prior distribution describes the experimentalist degree of belief for the true model parameters. An initial prior p_t=1_ (θ) was set in a discrete gridded parameter space which is comprised of five-dimensional vectors θ = (M, A, F, β, δ). The prior distribution for the parameters (M, A, F, β, δ) is a joint normal distribution ~N([2.00, -0.30, 0.78, 0.62, -0.30], diag(0.50, 0.50, 0.50, 0.11, 0.02)) in log10 space across a constrained 5-D parameter space, representing a weak prior knowledge of normal vision (Figure 2, cyan curve) and narrow priors on the “nuisance” parameters β, δ (Prins, 2013). Once the posterior distribution is updated, the simplest representation of its information is to assert a single point estimate of the parameter values. In the Bayesian framework, various methods provide the final point estimate: the mode of the posterior (MAP), the median of the posterior (MED), and the mean of the posterior (MEAN). Here we chose the MEAN as a final estimation of θ.

**Figure 2.**
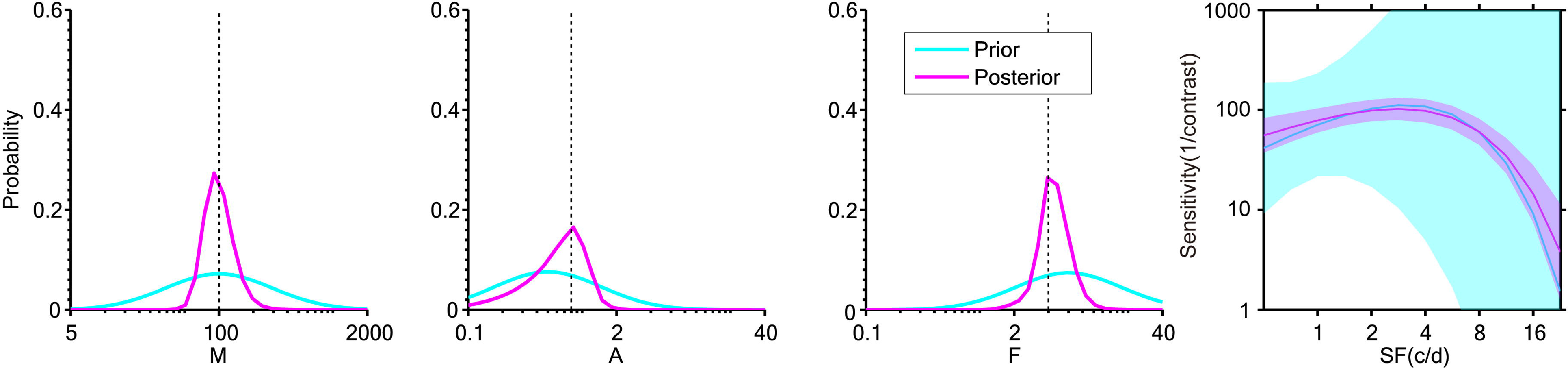
The figure illustrates the distribution of the three “key” parameters of CSF prior to the data collection (cyan) and after 108 2-AFC simulated trials (magenta). The rightmost plot shows the ±1 st.d. prior and posterior in the 2D CSF space. The dashed lines indicate the designated ‘true’ value in simulation. The plots show that: (1) The prior distribution represents a weak initial knowledge. (2) The posterior distribution after 108 2-AFC trials converges near the ‘true’ value of simulation.

### 1.3. Adaptive Methods

Four adaptive methods including two 2-D adaptive (FIG, qCSF) and two 1-D adaptive (Ψ, staircase) are described.

Fisher Information Gain (FIG). We implemented a maximum fisher information gain method (Liao & Carin, 2004; Remus & Collins, 2007) in which the sample was selected using a one-step-ahead search for maximizing the determinant of the Fisher information matrix. This method optimized sampling step to get maximum information about the parameters: Once the information gain had been calculated, the stimulus would be selected as the one corresponding to the maximum of gain of Fisher information. In practice, the stimulus value xt was selected uniformly among the top 10% of best stimuli, in order to avoid being trapped in local minima and to obtain a more uniform sampling of the SF space. With this constrain, the sampling of SF space was still biased toward the edge SFs (see Supplementary Material), because they are the most informative for the fitting procedure, but much less than without this constrain (results not shown). The prior probability distribution, p(θ) was generated over a 5-D parameter space, as described in the previous section Bayesian Inference, and also used for stimulus selection of the first trial. After the final trial, the parameters estimates were defined by the mean value across the whole parameter space and the estimated sensitivity was reported as FIG in the results.

qCSF. Quick contrast sensitivity function (qCSF) is a Bayesian adaptive procedure that was designed for concurrently estimating contrast thresholds across the full spatial-frequency range (L. A. Lesmes et al., 2010). Here, for the convenience of comparison with the other methods, we used our CSF model (Equation 2) and the prior distribution described in section Bayesian Inference, by changing the CSF in the qCSF toolbox and making a 2-D Bayesian re-estimate of sensitivities after each simulation (reported qCSF). As for the FIG, sampling of SF space was also biased to the “edge” SFs (see Suppl.Material), but somehow stronger.

The Ψ method (Kontsevich and Tyler, 1999) is a Bayesian adaptive technique that estimates the parameter values of a 1-D psychometric function from posterior distribution describing the combined distribution of parameters α-β. In our study, we used the Palamedes toolbox (Prins & Kingdom, 2009) and ran an independent Ψ measurement simultaneously and interleaved for each spatial frequency. The prior distribution of parameters α, β of every Ψ measurement was a joint normal distribution ~N([-log10(S(f)), 0.62], diag(5, 0.11)) in log10 space, where S(f) represented the prior sensitivity at the respective spatial frequency f. The prior distribution represented a weak prior knowledge of α and a constrained prior assumption of β. The reciprocal of the final threshold estimate of every Ψ run was the contrast sensitivity at the spatial frequency and was marked as Ψ in the article. The CSF was also estimated by the 2-D Bayesian inference and was marked as Bayes-Ψ in the article.

Up-Down Staircase. A simple up-down staircase method (3 up 1down) (Kaernbach, 1991) was used—the contrast stepped up ~33% after a negative response, and stepped down 10% after a positive response. For each spatial frequency an independent staircase was run and the staircase started from the prior estimated contrast threshold at the respective spatial frequency. To speed up the measurements, we made the first four trial 3-exponent step sizes (i.e. ~136% up and ~33% down). For every staircase, we averaged the contrasts of all trials to estimate the contrast threshold after excluding the first four trials. All these staircases were randomly interleaved. The CSF was then estimated by the 2-D Bayesian inference and was marked as Bayes-Stc in the article.

### 1.4. Simulation Methods

To investigate the efficiency of 2-D Bayesian inference in different adaptive strategies, Monte-Carlo simulations (N=1000) were used. We simulated 2-D Bayesian estimates from 4 sampling methods (FIG, qCSF, Bayes-Stc, and Bayes-Ψ) and a 1-D estimates (Ψ), as described in the previous sections. Each simulated experiment consisted of sampling trials M={48, 72, 108, 156, 228, 300}, spatial frequency ranging from 0.5 to 22.6 cycle/degree (cpd) in 0.5 log2 unit steps, and possible contrast values were from 0.001 to 1 in 0.02 (log10) unit steps. The initial stimulus point of every simulation was selected according to the sampling methods, which were described in details in section 1.3 (**Adaptive methods**). The simulated observer, specified by parameters {M, A, F, β, γ, δ} = {100, 0.8, 4, 4, 0.5, 0.02}, matches a normal observer in a 2AFC contrast detection task (Figure 1). For each simulated trial, these values were introduced into Equation 2 to generate the simulated response.

We also exemplified the influence of initial priors using data artificially generated from a curve poorly matching the prior, whose parameters were {M, A, F, β, γ, δ} = {40, 1.2, 6, 4, 0.5, 0.02}, representing an amblyope (C.-B. Huang et al., 2008). Demonstration codes in MATLAB (MathWorks, Natick, MA) and Octave (GNU) are available for download (http://vision.ustc.edu.cn/packages_en.html).

### 1.5. Experimental methods

#### 1.5.1. Apparatus

A vertically-oriented sinusoidal grating was displayed in the center of the screen (Sony MultiScan G520) driven by a nVidia Quadro K600 Engine with 500 MB of video RAM, housed in a Windows Intel Core 2 PC. A video switcher (Li, Lu, Xu, Jin, & Zhou, 2003) was used to generate a 14 bit gray level. The mean luminance of the screen was set to an absolute level of 48 cd/m^2^; the gamma function and parameters for the method were calibrated every day before the experiment, at least 30 minutes after monitor was switched on. The resolution was set to 1600*1200 at 85Hz. The display window was masked by a gray cardboard to a circle aperture subtending 4.2 deg at the usual viewing distance of 4m. To remove any sources of distraction all data collection took place in a dark room. The stimuli were monocularly viewed by subjects’ dominant eyes with the fellow eye patched.

#### 1.5.2. Stimuli

Vertically oriented sinusoidal gratings were presented in a 3 degree circular window. A Gaussian cumulative function distribution was used to blend the grating’s edge into the background. Every stimulus grating had a random phase. The grating was presented with a limited-lifetime of 150ms in every interval.

#### 1.5.3. Subjects

The subjects (5 naive observers; age: 23 to 30 years; 2 males) had normal or corrected-to-normal vision and were experienced at the task. Written informed consent was obtained from the subjects after explanation of the nature and possible consequences of the study. The experiments were conducted according to the experimental protocol for human subjects approved by the ethics committee (IRB) of the School of Life Science, University of Science and Technology of China.

#### 1.5.4. Procedure

Subjects were seated in a dimly lit room and head stabilized with a chin-rest. Observers were presented with a 2-interval forced choice (2-IFC) task. On each trial, two intervals separated by a 500 ms gap were presented for 150 ms. In one of the intervals the target grating was presented, and in the other the mean luminance background stayed on. The observers’ task was to indicate by keyboard pressing that in which interval the target grating appeared. An intermediate frequency pure tone was provided at the beginning of every interval and a high frequency pure tone was provided after every response, irrespective of response correctness.

All participants completed a series of CSF runs. Two adaptive methods were used, the Ψ method and FIG method. For both methods, spatial frequency values were 0.5 to 22.6 cycle/degree (cpd) in 0.5 log2 unit steps, and possible contrast values were from 0.001 to 1 in 0.02 log10 unit steps. The initial stimulus point of every measurement was selected according to the sampling methods, which were described in details in section 1.3, Adaptive methods.Each measurement contained a total of 108 trials, and all stimuli and responses were used in the Bayesian inference of the CSF. A session was constructed by 2 methods with 3 repetitions per method, providing a total of 648 trials per session (or day). All these 648 trials were interleaved. The subjects were tested over 4 days, i.e. 2592 trials, totally. Subjects were preliminary trained for 216 trials one day ahead of the experiment and 24 practice trials every day before the experiment (Jakel & Wichmann, 2006). These practice trials were not included in the results and analysis. We further performed a 2-D Bayesian inference with the data sets sampled by the Ψ method, such that we obtained three estimated CSF values (FIG, Bayes-Ψ, and Ψ) for every subject.

## 2. Results

### 2.1. Simulation

In the experiment, we analyzed the efficiencies of 2 kinds of fancy 2-D adaptive estimates (FIG and qCSF), 2 types of 2-D Bayesian estimates with usual strategies (Bayes-Ψ and Bayes-Stc), and two usual estimates (Ψ from 1D fitting, and Stc from 1D convergence point estimates). Figure 3 shows the 6 CSFs estimated after 108 sampling trials. The error region (shaded) represents the variability (mean ± 1 st.d.) for estimating individual thresholds at a given spatial frequency. Out of the Ψ and Stc methods, it seemed that all other sampling schemes provided very similar results and distributions. To quantify the concordance of CSF estimates, we calculated the root mean squared error (RMSE) of the threshold obtained from the 6 methods, collapsed across all simulations (N=1000) and spatial frequency conditions (S=12) (see also Hou et al. 2010). As shown in Figure 4, the RMSE of sensitivities estimated with the FIG, qCSF, Bayes-Ψ, Bayes-Stc, Ψ, and Stc at 108 trials were 1.9, 2.2, 2.1, 2.2, 4.9, and 4.7 dB respectively, and all decreased as the trial number increases. The results showed that the 2-D Bayesian inference has a considerable effect of decreasing the estimated RMSEs of usual estimates (1-D estimates), and, for example, the average RMSEs of Ψ among trials decrease from 4.9 dB to 2.2 dB with the help of 2-D estimates. In other words, the Bayes-Ψ yielded a relatively large advantage compared to Ψ, obtaining the same precision within almost one fourth trial numbers: 168 trials of Ψ provided precisions of about 48 trials of Bayes-Ψ, and 228 trials of Ψ correspond to about 72 trials of Bayes-Ψ. The improvements of efficiency of another usual method, the staircase, was similar.

**Figure 3.**
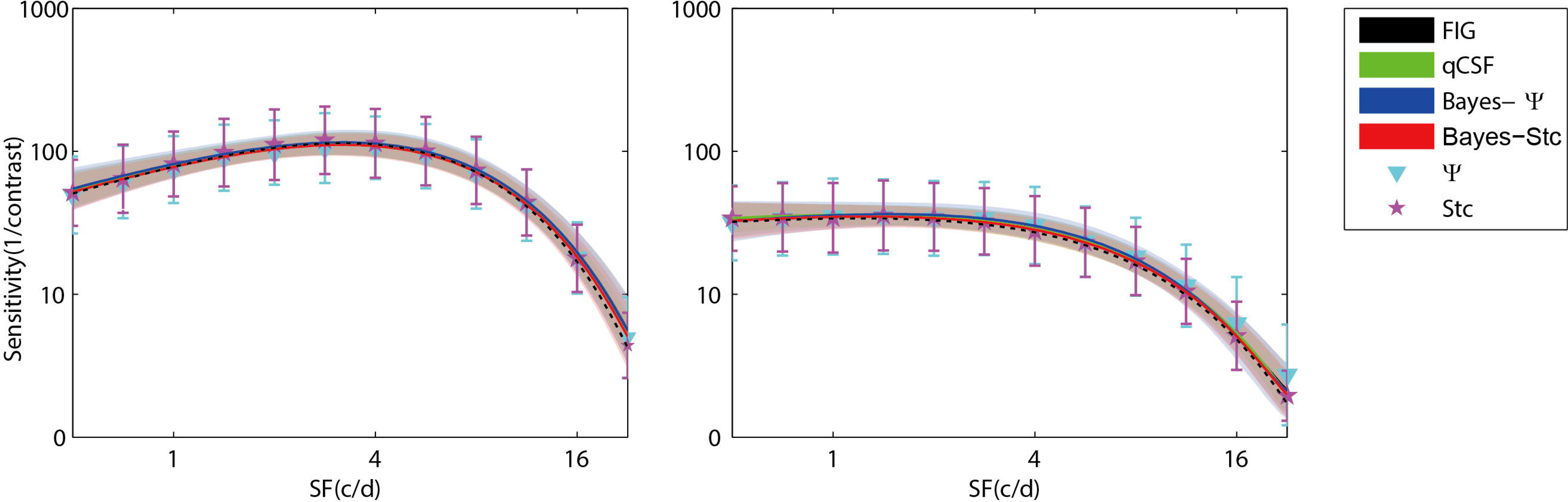
Results of simulating a normal observer (left) and an amblyopic observer (right). CSFs obtained with FIG, qCSF, Bayes-Ψ, Bayes-Stc, Ψ, and Stc methods for 108 trials. The shaded region and error bar represent 1 st.d.

We also analyzed the accuracy defined as the bias to the ‘true’ threshold value in dB unit. Figure 4 depicts the bias for threshold estimation with each of the five methods. These 6 estimates exhibited similar small biases, which were 0.48, 0.35, 0.45, 0.14, -0.13, and 0.33 dB after 108 trials, respectively for the FIG, qCSF, Bayes-Ψ, Bayes-Stc, Ψ, and Stc. The absolute value of all biases except Stc’s decreased as trial number increased.

**Figure 4.**
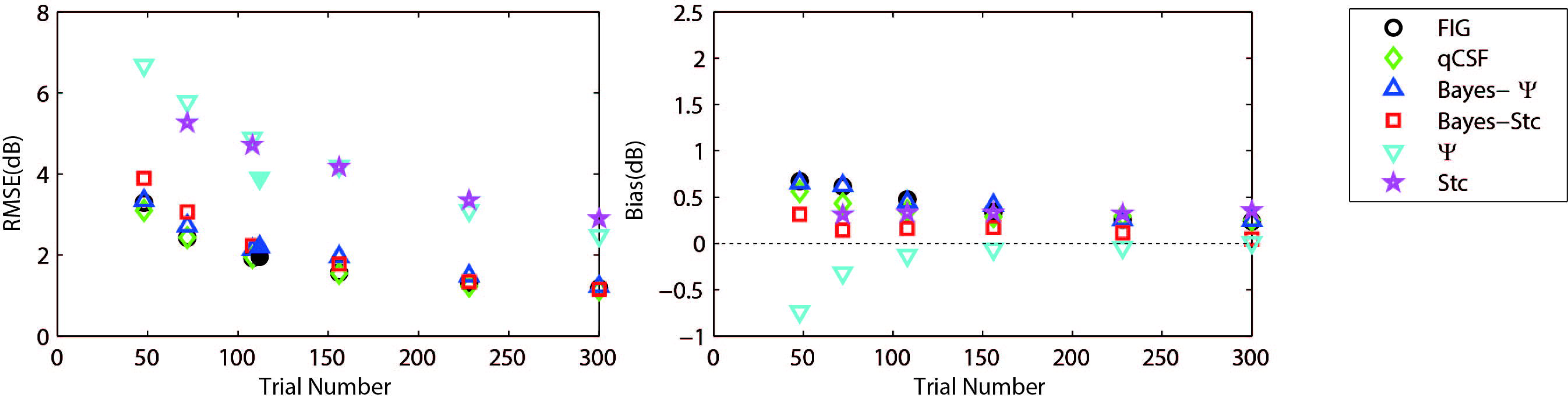
The average precision (left) and relative bias (right) estimates of the methods in simulation (open symbols) and psychophysical experiments (filled symbols). The symbols of the experimental results are slightly shifted for clarity purpose. Different methods are represented by different colors and symbols (see legend).

To further demonstrate that the test efficiencies exhibited by the current simulation were not overly determined by the initial priors, we simulated the Bayesian measurement of a widely different CSF (Figure 4 right) observed for an amblyope (C.-B. Huang et al., 2008). As shown in Figure 5, CSF estimates provided by the Bayesian methods converged to 2.06, 1.79, 2.05, 2.01, 5.56, and 4.66 dB for measuring methods, FIG, qCSF, Bayes-Ψ, Bayes-Stc, Ψ, and Stc, by the 108^th^ trial; the mean bias magnitude, 0.63, 0.63, 0.63, 0.68, 0.99, and 0.46 dB, continued to decrease with more trials. These results are comparable to the results above of a CSF close to the prior peak.

**Figure 5.**
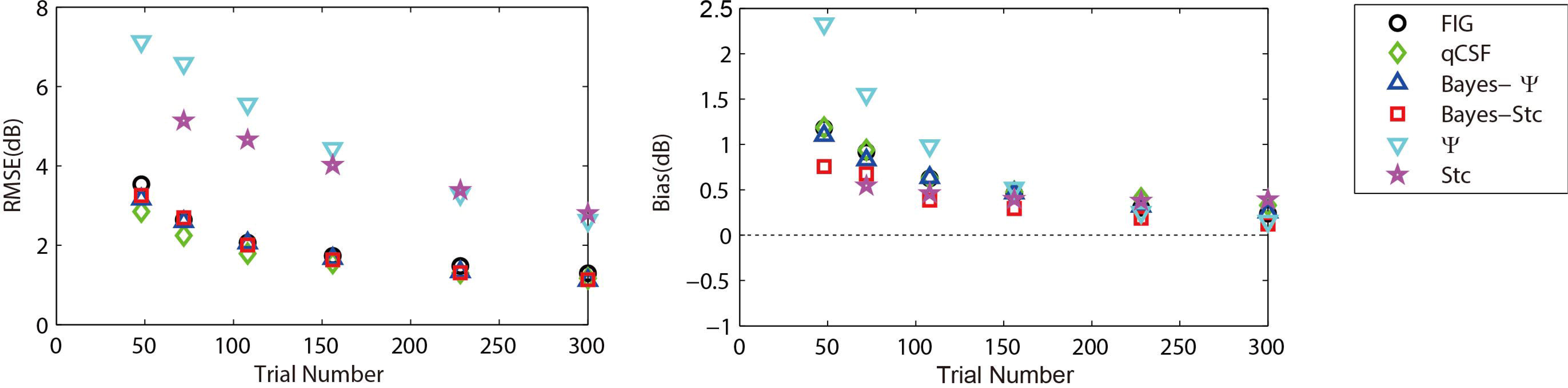
The average precision (left) and relative bias (right) estimates of the methods in simulating an amblyopic observer. Different methods are represented by different colors and symbols (see legend).

### 2.2. Experimental test

A standard 2-interval-forced-choice grating detection task with 108 trials was used for psychophysical validation of the Bayesian framework. A fancy 2-D adaptive method, FIG, and a traditional sampling method, Ψ, were applied independently and repeatedly for each of the 5 subjects. We evaluated the precision of the methods through RMSE of repeated measurements across four days and 3 repetitions per day and subject.

The measured CSFs efficiency of the 5 subjects for the three estimates (Bayesian-FIG, Bayes-Ψ, and Ψ) are plotted on Figure 6. The errors (shaded region or error bar) represents the variability (± 1 st.d.) for estimating individual thresholds. To quantify the concordance of CSF estimates, we computed the RMSE of the threshold obtained with the three methods, collapsed across all observers (O=5), repetitions (N=12) and spatial frequency conditions (S=12). The results were added to the simulations in Figure 4. The RMSE estimated with the FIG, Bayes-Ψ, and Ψ methods were 2.0, 2.2, and 3.9 dB respectively (Figure 4, filled symbols). The 2-D Bayesian inference has a considerable effect of decreasing the estimated variance—the RMSE of Ψ estimates decrease from 3.9 dB to 2.2 dB after a Bayesian inference. And the precision was comparable with the fancy estimate (FIG), with the difference being only 0.2 dB.

**Figure 6.**
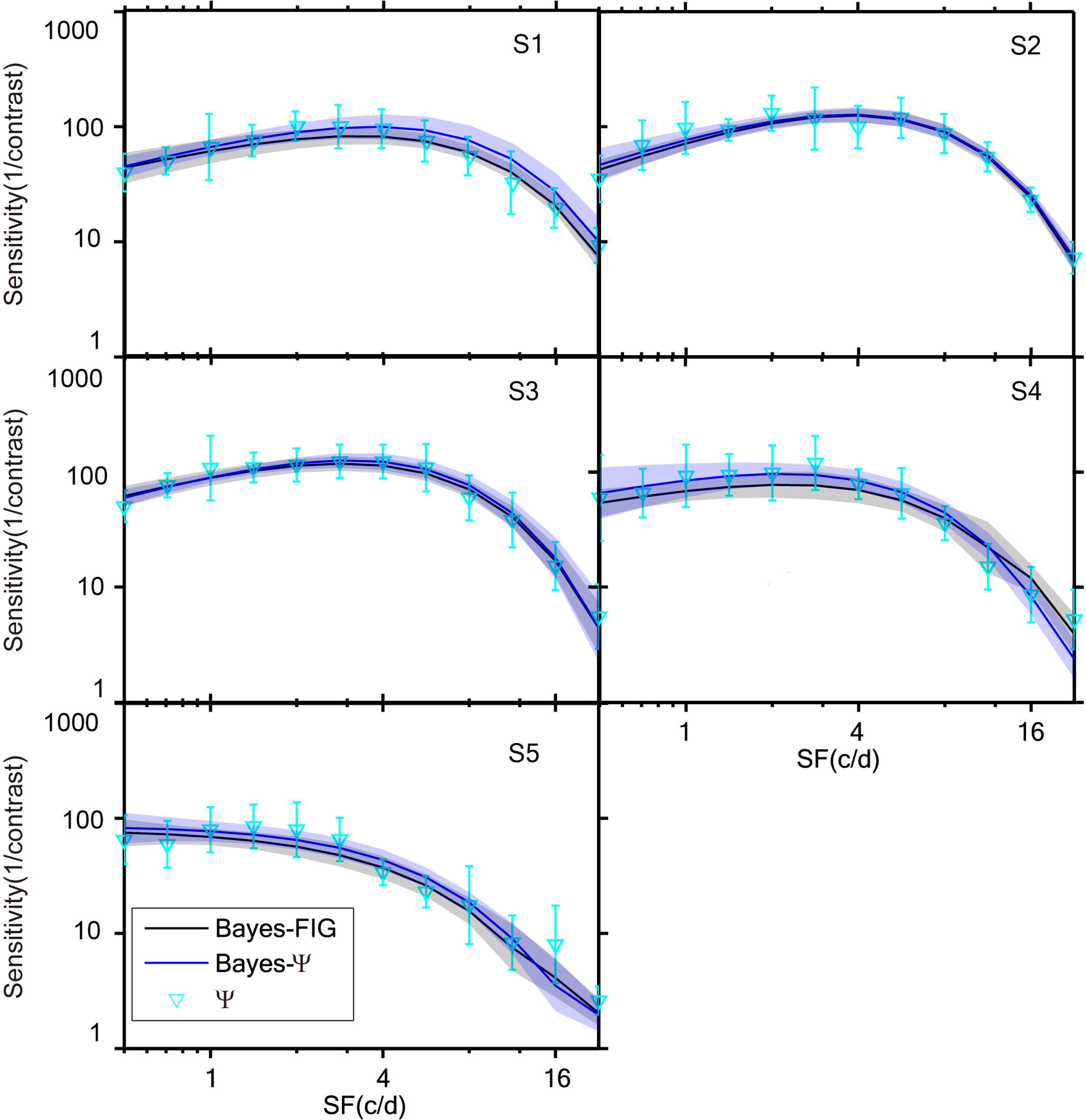
CSFs measured for 5 subjects with FIG, Bayes-Ψ, and Ψ methods repeated 12 times with 108 trials. Both Bayesian methods have smaller variability than the traditional method. The shaded region and error bar represent 1 st.d.

## 3. Discussion

Our study demonstrated that the 2-D Bayesian framework strongly increases the efficiency of traditional simple adaptive methods and makes them comparable with more recent fancy 2-D parametric adaptive methods. We performed a large number of simulations and showed that this framework was fast and convenient and could be applied to behavioral data sets sampled by usual simple strategies. This method presented an acceptable compromise that allowed for efficient estimation of CSF with traditional simple adaptive strategies. Under a psychophysical validation, the method improved the accuracy in a similar way as in the simulations, and showed a good applicability in real conditions.

The existing sampling procedures considered in this study (FIG, qCSF, Stc, Ψ) are four examples of the numerous adaptive techniques available for psychometric testing. The staircase method (Stc), simplest of all sampling methods, requires very few assumptions and have fairly simple algorithms for placement of stimuli. There is another general category of adaptive methods, parametric adaptive methods, which sample a stimulus by applying complicated algorithms on estimates from previous samples. These parametric ‘sample-estimate-sample’ strategies thus make a positive-feedback loop, and face a risk of trapping in local minima (L. A. Lesmes et al., 2010). Fortunately, researchers have already invented numerous toolboxes (L. A. Lesmes et al., 2010; Prins & Kingdom, 2009; Shen, Dai, & Richards, 2015) to help with simplifying the application and escaping from local minima. They also proved the methods’ robustness in estimating anomalous functions (Hou et al., 2010; Luis Andres Lesmes, Jackson, & Bex, 2013). However, there still exist the concerns of complexity and potential traps of these fancy 2-D parametric adaptive methods. And the usual methods are still widely accepted by psychophysical researchers for their robustness and simplicity in applications (Bonneh et al., 2016; Chung & Legge, 2016; Klein, 2001; Richard et al., 2015; Vedamurthy et al., 2015). Our 2-D Bayesian framework improve the efficiency of traditional methods and provide researchers with a flexible choice.

Bayesian inference approach is often criticized for its dependence on prior, but it also provides a straight-forward and reasonable way to realize constraints of function parameters (Kuss et al., 2005). We have chosen an almost flat prior distribution across a wide magnitudes of CSF parameters (Figure 2) to avoid any mis-predefinition of the parameters. The robustness of the 2-D Bayesian framework was demonstrated by estimating a CSF that poorly matched the prior curve (Figure 6).

The parameters describing the slope, guessing rate, and lapsing rate are considered to be nuisance parameters, since the parameters do not describe the interested sensory mechanism but nevertheless do affect our observations. Wichmann and Hill (2001) have shown that the threshold and slope estimates of a psychometric function may be severely biased when it is assumed that the lapse rate equals zero but lapses do, in fact, occur. In the Bayesian framework, as Prins (2013) nicely demonstrated, we can give the nuisance parameters proper attention and propose a strategy that limits the prior guess and lapse rate in a narrow normal distribution. This method provides small bias changes in parameter estimates in our application too (see Supplementary Material).

To summarize, the 2-D Bayesian inference framework appears to be a good choice for estimating the parameters of the contrast sensitivity function applicable to any sampling strategy. Besides the four adaptive strategies considered in the study, the framework should be also applicable to other strategies. Furthermore, it is flexible and could be applied to measure other behavioral functions that links subjects’ binomial-distributed responses to multi-dimensional stimulus spaces (e.g. color discrimination in a three-dimensional RGB color space, motion contrast sensitivity in a speed-contrast space, or any other psychophysical function). Applying the method provides the experimenter with the freedom to use a stimulus sampling procedure appropriate to their research interest and experience, while still estimating the interested function in a highly efficient way.

